# Niche preference of *Escherichia coli* in a peri-urban pond ecosystem

**DOI:** 10.1101/2020.01.30.926667

**Authors:** Gitanjali NandaKafle, Taylor Huegen, Sarah C. Potgieter, Emma Steenkamp, Stephanus N. Venter, Volker S. Brözel

## Abstract

*Escherichia coli* comprises of diverse strains with a large accessory genome, indicating functional diversity and the ability to adapt to a range of niches. Specific strains would display greatest fitness in niches matching their combination of phenotypic traits. Given this hypothesis, we sought to determine whether *E. coli* in a peri-urban pond and associated cattle pasture display niche preference. Samples were collected from water, sediment, aquatic plants, water snails associated with the pond as well as bovine feces from cattle in an adjacent pasture. Isolates (120) were obtained after plating on Membrane Lactose Glucuronide Agar (MLGA). We used the *uidA* and *mutS* sequences for all isolates to determine phylogeny by maximum likelihood, and population structure through gene flow analysis. PCR was used to allocate isolates to phylogroups and to determine the presence of pathogenicity / virulence genes (*stxI, stxII, eaeA, hlyA*, ST and LT). Antimicrobial resistance was determined using a disk diffusion assay for Tetracycline, Gentamicin, Ciprofloxacin, Meropenem, Ceftriaxone, and Azitrhomycin. Our results showed that isolates from water, sediment and water plants were similar by phylogroup distribution, virulence gene distribution and antibiotic resistance while both snail and feces populations were significantly different. Few of the feces isolates were significantly similar to aquatic ones, and most of the snail isolates were also different. Population structure analysis indicated three genetic backgrounds associated with bovine, snail and aquatic environments. Collectively these data support niche preference of *E. coli* isolates occurring in this ecosystem.

## Introduction

*Escherichia coli* is a commensal in the gastrointestinal tracts of humans and vertebrate animals, but readily isolated from aquatic and terrestrial habitats. Some data suggest semi-permanent residence in extra-host habitats [1, 2]. The species displays a broad range of genotypes and associated phenotypes [3, 4] and has been classified into four phylogroups (A, B1, B2 and D) [5], and later eight phylogroups based on their genomic information [6]. Of these, seven (A, B1, B2, C, D, E and F) belong to *E. coli sensu stricto* whereas the eighth is represented by cryptic Clade-I. Variation in genotype and phenotype among strains of different phylogroups is believed to support fitness in different ecological habitats, leading to niche preference. Phylogroups A and B1 occur more frequently in the environment [7]. Some strains of phylogroup B1 were reported to persist in water [7, 8] and soil [9], with some believed to be naturalized members of their specific communities [1]. B2 and D strains are frequently isolated from extra-intestinal sites within host bodies [3]. Many studies have reported that phylogroup B2 and, to a lesser extent D strains are more likely to carry virulence factors than other phylogroups [10-12]. Interestingly, virulence genes are more frequently present in phylogroup B1 isolates from environments where phylogroup B2 strains are absent [13]. Thus, identification of the phylogroup of unknown isolates may provide information on their physiological characteristics and ecological preferences.

*E. coli* is a highly diverse species as revealed by DNA fingerprinting of populations obtained from different sources [14-17]. Genomic analyses also showed that *E. coli* has a large open pan-genome (Rasko et al. 2008; Touchon et al. 2009) with an estimated core genome of less than 1,500 genes but an accessory genome containing a reservoir of more than 22,000 genes (Robins-Browne et al. 2016). Of the latter many may be uncharacterized yet important virulence factors [18]. The pan-genome of 61 *E. coli* comprised 15,741 gene families and only 993 (65) of these families were represented in every genome, comprising the core genome [19]. Similarly, genome data for 228 *E. coli* isolates revealed a pangenome of 11,401 genes of which 2,722 (23.9%) were core [20]. The large genomic and resulting phentotypic diversity explains the versatile behavior of this species. It exhibits a biphasic life style with primary habitat in the mammalian gastrointestinal tract, but water, sediments and soils are reported as non-host or secondary habitat [7, 21-24]. Extensive genetic diversity was also reported for the *E. coli* populations within different environmental habitats [25-27]. Several studies further reported that *E. coli* survive and grow in water, sediments, soil, and on water-plants in various climatic regions where no evidence for fecal contamination exists [21, 28-30]. Certain strains of *E. coli* present in the environment also appear to have become naturalized, and to have distinct genotypes when compared to strains associated with animal hosts [9, 21]. Previous studies have suggested that there is a relationship between genotypes of *E. coli* found among specific animal hosts and the geographic location from which they were isolated [21, 30, 31]. The genetic differences between populations are a consequence of continued evolutionary success due to their survival and adaptation in different environments [32]. This suggests that *E. coli* strains would perform best in niches that match their specific combination of phenotypic traits.

We hypothesized that *E. coli* display niche preference when presented with multiple environments, and that some strains would display fitness in aquatic environments over the mammalian gut. A secluded peri-urban pond adjacent to a cattle pasture was selected as sampling site. We isolated *E. coli* from the water, sediment, submerged water plants and water snails, as well as from bovine feces in the adjacent pasture. To determine evidence of niche partitioning, isolates were characterized genotypically by phylogrouping, analysis of their *uidA* and *mutS* sequences, and virulence gene distribution, and phenotypically for antibiotic resistance.

## Materials and Methods

### Sample source

Samples were collected from a secluded pond (GPS co-ordinate 44.2719° N, 96.7736° W) at the edge of Brookings, SD, USA during June and July 2013. This pond is located between the edge of town, a nature park, and a cattle pasture, and surrounded by dense scrub and trees, rarely visited by humans. Water (31), sediment (27), water plant (35), and snail samples (20) were collected from the pond, and bovine feces (7) was collected from the adjoining cattle pasture. Samples were placed into sterile 50 mL conical screw cap tubes, brought to the laboratory on ice and processed on the same day.

### Isolation of *E. coli*

Water samples (10 mL and 1 mL) were filtered through a sterile 0.45 μm mixed cellulose ester filter (Milipore) and the filters placed on Membrane Lactose Glucuronide agar (MLGA, Fluka analytical). Sediment samples were mixed with 15 mL of sterile dH_2_O, shaken for 30 s, and 1 and 10 mL aliquots filtered before placing filters onto MLGA. Water plants and snails were rinsed with sterile dH_2_O, then crushed in 10 mL sterile dH_2_O, and 100 μL plated directly on to MLGA plates. Feces samples were suspended in sterile water and serial tenfold dilutions plated onto MLGA. MLGA plates were incubated at 37°C for 18h. Green colonies indicated positive for β-Galactosidase (yellow) and β-Glucuronidase (blue), and were assumed to be *E. coli*. This protocol therefore excluded ß-glucuronidase negative O157:H7 strains. One colony was selected at random from the highest dilution showing growth, streaked onto MLGA to confirm purity, sub-cultured on LB agar, and stored at -80°C in 50% glycerol.

### Phylogroup analysis

Genomic DNA was extracted from overnight LB agar cultures harvested and re-suspended in 5mL 10 mM phosphate buffer (pH 7.0) using the genomic DNA Quick Prep Kit (Zymo Research), and stored at -20°C. Isolates were assigned to phylogroups using the protocol of Clermont, Christenson (6). To avoid ambiguity, PCR was performed separately for each primer set (Table S1). Phylogroup similarity among the five sample types was determined by UPGMA analysis using the constrained Jaccard coefficient in PAST version 3.14 (http://folk.uio.no/ohammer/past) [33]. To determine whether the distribution of phylogroups differed by source we used multinomial log-linear regression models. The models were fitted using the nnet package in R (v.3.2.2)[34]. The response variable in this analysis was the phylogroup of each isolate (A, B1, B2, C, D, E, and Unknown), and the explanatory variables were the sample source and clusters associated with origin of the isolates. To visualize the effect of significant explanatory variables, we used regression trees fitted using Package Party [35] in R.

### *uidA* and *mutS* sequence analysis

The *uidA* and *mutS* genes were amplified by PCR using primers described previously [36] (Table S1). PCR reactions (25 µl) were set up as follows: 2.5 µl reaction buffer (10X) (New England Biolab), 1.5 µl MgCl_2_ (25mM), 0.5 µl dNTPs (40mM), 0.1 µl forward primer and 0.1 µl reverse primer (100 µmol), 0.125 µl of Taq polymerase (NE Biolabs), 0.5 µl of DNA template and 20.7 µl sterile nano pure water. The amplification cycle was initiated with 95°C for 2 min, followed by 30 cycles of denaturing at 95°C for 30 s, annealing at 56°C for 30 s and extension at 72°C for 1 min, with a final extension at 72°C for 5 min. DNA sequences were determined by the dideoxy chain termination method (Beckman Coulter Genomic Center at Denver, MA). The *uidA* and *mutS* sequences were submitted to Genbank (http://www.ncbi.nlm.nih.gov/genbank/) under BankIt2031081: MF459726 - MF459846 and BankIt2031086: MF459847 - MF459967 respectively.

To infer the relationships among isolates, DNA sequences were aligned using ClustalW [37], and overhangs were trimmed using SeAl [38]. The *uidA* and *mutS* sequences for all isolates and reference strains [39] were concatenated using SeAl. A maximum likelihood analysis using model GTR+G+I with 1,000 bootstrap replicates was performed in the program MEGA 6.06 [37]. The tree was then annotated and visualized using the ITOL online tool [40].

### Population Genetic Analysis

To infer population structure and assign isolates to distinct populations, we employed a model-based clustering method using STRUCTURE [41]. More specifically the admixture model was applied using sample locations as prior (LOCPRIOR). By assuming mixed ancestry, individuals within a population were thought to have inherited a fraction of their genome from an ancestor in the population [42]. Ln probability values and the variance of Ln likelihood scores were estimated for the concatenated *uidA-mutS* sequences, assuming the presence of 2 populations (K = 2, with an adjusted alpha = 0.5) and performing twenty iterations for each K from K = 1 to K = 6. For these analyses a burn-in of 10,000 and a run length of 500,000 were used [43]. All other parameters in STRUCTURE were left as default. The resulting data from STRUCTURE were collated and visualised using the web-based program Structure Harvester [42] to assess which likelihood values across the multiple estimates of K best explained the data (in this case K=3 was the best) using the Evanno method [44, 45]. Furthermore, optimal alignments for the number of replicate cluster analyses were generated using the FullSearch algorithm in CLUMPP [46] and the corresponding output files were used directly for cluster visualization as plots in Excel and the program Distruct 1.1 [47].

### Virulence gene assays

PCR for detection of *stx1*, stx2, *eaeA* and *hlyA* genes was performed using primers as described by Fagan, Hornitzky (48) (Table S-2), and for ST and LT virulence genes as described by Osek (49) (Table-S2). DNA samples for PCR were prepared by the boiling method. Stock cultures were recovered on LBA, two colonies suspended in 500 uL dH_2_O, washed by centrifugation and suspended in sterile dH_2_O, lysed by incubating at 100°C for 10 min, and immediately chilled on ice for 5 min. Debris was removed by centrifugation for 1 min at 12,000 X g and the supernatant was transferred to a new sterile tube and stored at -20°C for further use as PCR template. PCR reactions were carried out in 25 µl volume containing 1 µl of DNA template, 2.5 µl reaction buffer (10X) (New England Biolabs), 1.5 µl MgCl_2_ (25mM), 0.5 µl dNTPs (40mM), 0.1 µl forward primer and 0.1 µl reverse primer (100 µmol), 0.1 µl of Taq polymerase (New England Biolabs), and 19.2 µl sterile nano pure water. PCR amplification for *stx1, stx2, eaeA*, and *hlyA* was performed under the following conditions: initial 95°C denaturation step for 3 min followed by 35 cycles of 20 s denaturation at 95°C, 40 s primer annealing at 58°C, and 90 s extension at 72°C. The final cycle was followed by a 72°C incubation for 5 min [48]. LT and ST were amplified under the following conditions: an initial DNA denaturation step at 94 C for 5 min followed by 30 cycles of 1 min of denaturation at 94°C, 1 min of primer annealing at 55°C, and 2 min of extension at 72°C. The final extension step was performed at 72°C for 5 min [49].

### Antibiotic resistance assays

Antibiotic susceptibility of the 120 *E. coli* isolates was determined using a disk diffusion assay following the CLSI standard [50]. Stock cultures were recovered in 5 mL Mueller Hinton (MH, Oxoid) broth at 37°C for 16h. Cells were harvested by centrifugation (10,000 X g, 2 min), re-suspended in sterile tap water and the cell density adjusted to 0.5 on the McFarland turbidity standard. Cell suspensions were spread onto MH agar (Oxoid), and antibiotic disks (Oxoid) (Ciprofloxacin (CIP, 5 μg), Meropenem (MEM, 10 μg), Ceftriaxone (CRO, 30 μg), Gentamicin (CN, 10 μg), Azithromycin (AZM, 15 μg), Tetracycline (TE, 30 μg), with Penicillin (10 μg) as control were placed on the surface. After 18h incubation at 37°C, zone diameters were measured and isolates scored as intermediately or fully resistant. Isolates resistant to two or more antimicrobials were defined as multidrug resistant. *E. coli* ATCC 25922 was included in each assay as a negative control as it is sensitive to all these antibiotics.

## Results

### Bacterial isolates

Strains of *E. coli* were obtained from water (31), sediment (27), water plants (35), and snails collected from the pond (20), and from fresh bovine feces (7) obtained from the adjacent pasture.

### Phylogroup Distribution

Isolate collections obtained from the water and submerged water plants showed similar phylogroup distribution (Fig. 1), predominated by phylogroups B1, E and some B2 isolates. Sediment was similar to water and water plants, but with the addition of phylogroup A strains. In contrast, snail isolates were mostly phylogroup B2, while those from bovine feces were phylogroup E. Multinomial log linear regression supported a significant difference (p < 0.001) between the isolate collections from water snails and those from water, sediment and plants (Fig. 1S), with the latter three collections not significantly different from one another. As our isolation method was based on MLGA (β-Glucuronidase and β-Galactosidase), phylogroup E strains lacking the *uidA* gene for β-Glucuronidase would have been excluded [51]. Yet, we obtained several isolates from feces, all phylogroup E.

**Fig. 1.**
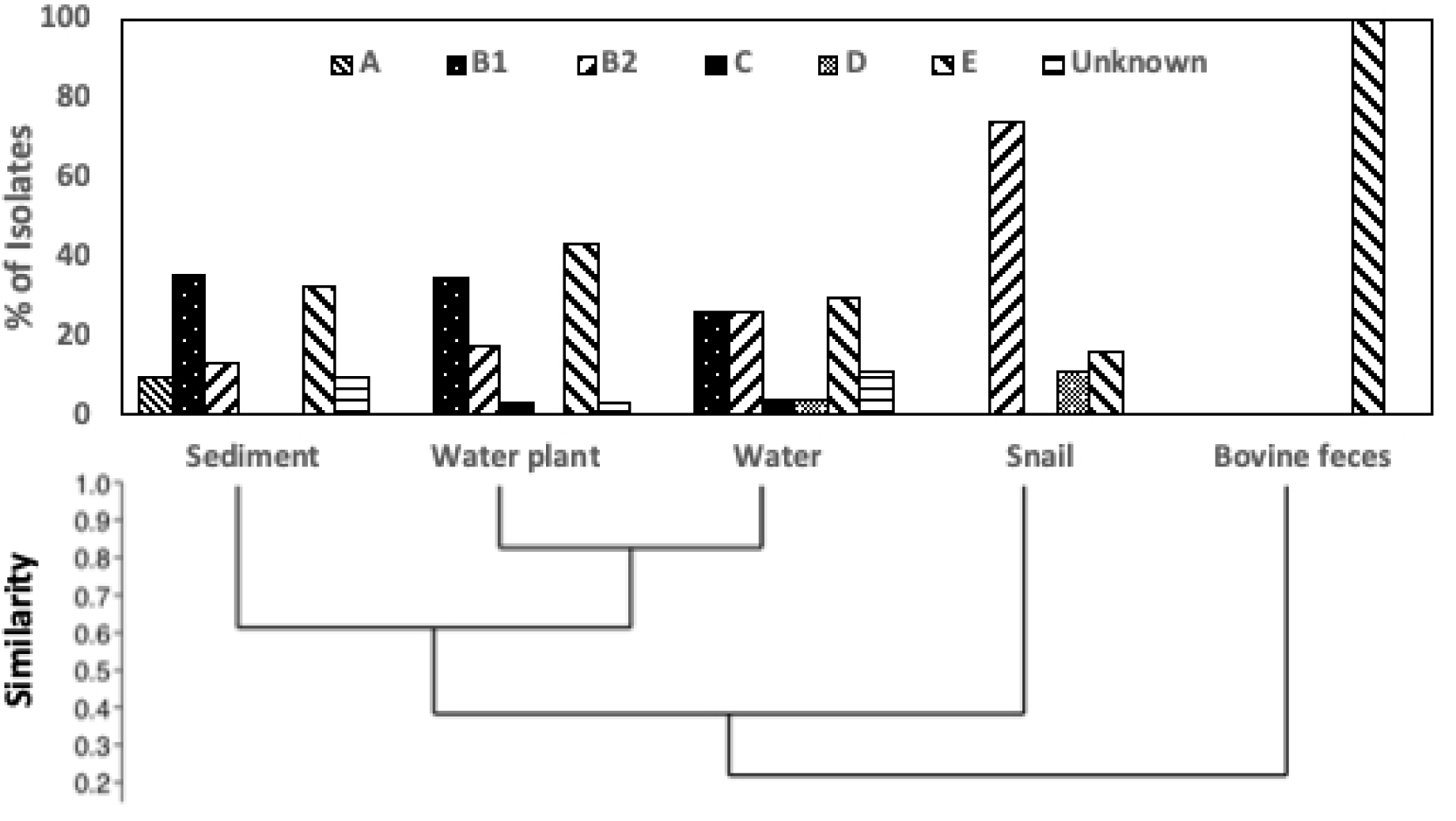
Phylogroup distribution across isolates from the five sample types. Phylogrouping was performed according to the scheme of Clermont et al., 2013. The relatedness between Phylogroup distribution similarity was determined by UPGMA using the constrained Jaccard coefficient.

### Phylogenetic analysis

The concatenated *mutS* and *uidA* sequence phylogeny formed many well-separated clusters with strong bootstrap support (Fig. 2). None of our isolates grouped with any of the Clade I, III, IV or V strains and all belonged to *E. coli sensu stricto*. Most of the water, water plant, and sediment isolates fell into mixed clusters, some with reference strains. This indicated co-occurrence of diverse strains across the three niches. The majority of water snail isolates grouped into three unique clusters that contained no water, sediment or water plant isolates, and also no reference strains, indicating that they are unique and potentially have a preference for snails over surrounding water, sediment or water-niches. Three of the snail isolates did cluster with water plant, sediment, water and reference strains. All bovine fecal isolates formed a separate cluster from aquatic and from snail isolates. Furthermore, no bovine isolates clustered with phylogroup E reference strains (Fig. 2), indicating hitherto poorly studied diversity within cattle.

**Fig. 2.**
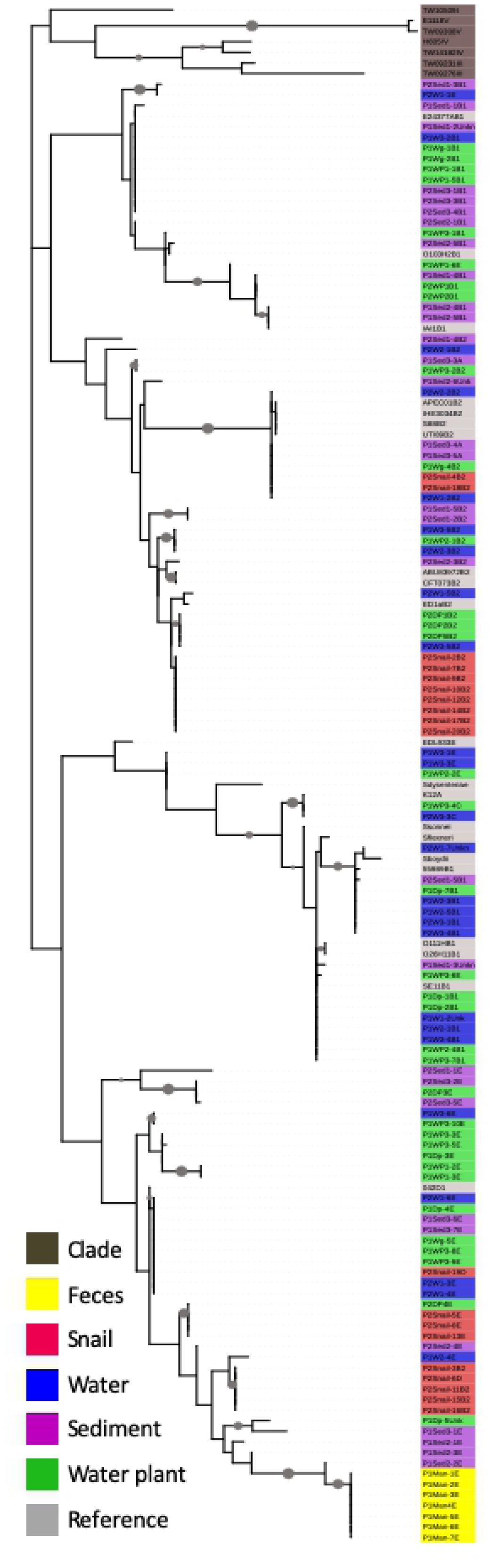
Phylogenetic analysis of the concatenated *uidA* and *mutS* gene sequences of *E. coli* isolates, reference strains and cryptic species of *E. coli*. Sequences were aligned using ClustalW and manually trimmed using Se-Al. The best Model: Maximum Likelihood analysis with GTR and G+I was performed in the program MEGA 6. The phylogenetic tree was color-coded and visualized using the Interactive Tree of Life with isolates color-coded based on their sources. Grey circles on branches indicate a bootstrap value of > 80% (1000 bootstraps).

### Population Genetic Analysis

Population genetic analysis of concatenated *uidA* and *mutS* genes was performed assuming one aquatic and one fecal population (i.e. K = 2, alpha = 0.5). The result obtained from the Evanno table was K = 3, supporting the existence of three separate genetic backgrounds within the collection of isolates examined (Fig. 3). The bovine fecal isolate collection was homogenous, containing mainly one genetic background. Isolates from snails were associated with two backgrounds that were mostly homogenous, one of which was identical to the fecal background. In contrast, water, sediment and water plant isolates were associated with a mixture of three genetic backgrounds shared by the bovine fecal isoloates, some shared by the second group of snail isolates, and a distinct third background (yellow in Fig. 3) that was more common in the aquatic populations but not in snail isolates. Thus the pond ecosystem comprised of an admixture of strains representing three genetic backgrounds, one likely due to introduction of bovine-derived strains (blue in Fig. 3), a second associated with snail populations (red), and a third unique to the aquatic environment (yellow). This indicates gene flow among the water, sediment and water plant populations, but with some genetic input from the fecal and snail populations. No genetic input from the water, water plant, sediment, and snail populations to the bovine fecal population was observed.

**Fig. 3.**
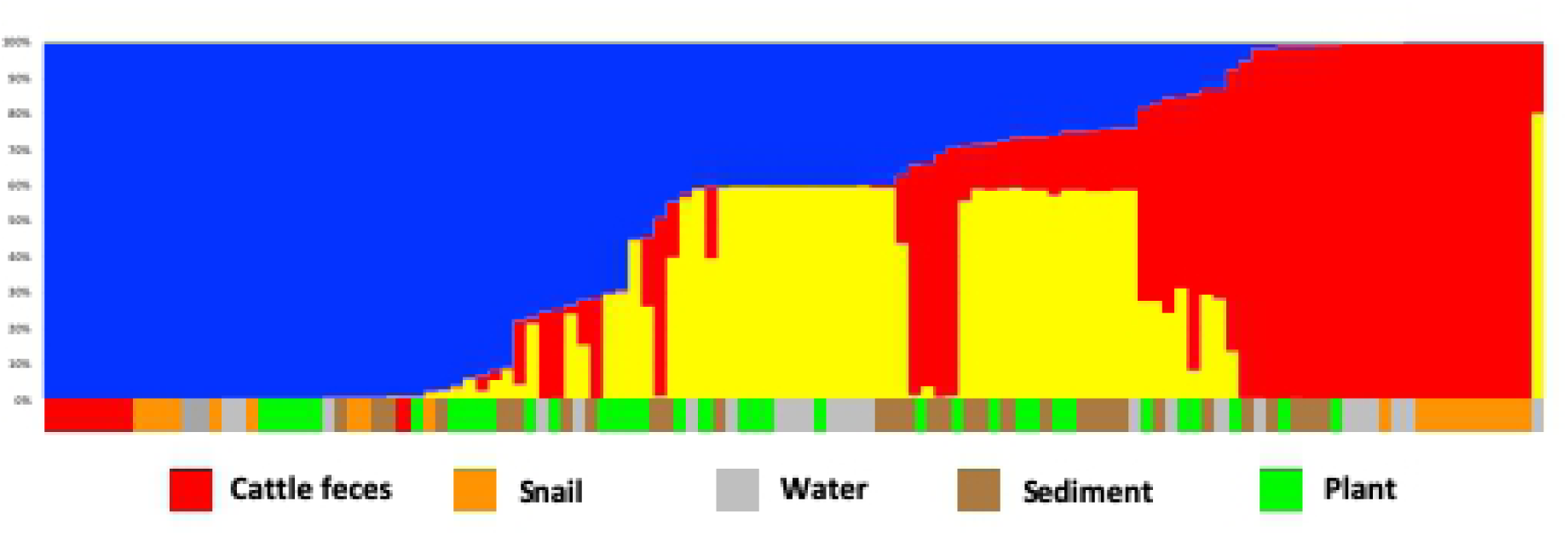
Population structure analysis of E. coli isolates. Concatenated uidA and mutS sequences were analyzed assuming presence of two populations, but analylis using Structure Harvester showed that K=3 best explained the data. The short color bars below the figure indicate the isolate source as defined in the legend.

### Virulence Gene Distribution

To determine their pathogenic potential, isolates were screened for the presence of major virulence genes associated with diarrhaeagenic *E. coli*. Out of six genes, four (*stx2, eaeA, hlyA* and *STb*) were detected. We did not detect any isolates with the *Stx-1* and LT genes, although the control strains EDL933D and O157:K88 [52] yielded positive results, confirming reliability of the assay. Among the four genes present, *eaeA* was the most frequently detected (36.13%), then *Stx2* (12. 61%), LTa (10.9%) and *hlyA* (3.36%). Distribution of the virulence genes in *E. coli* populations of water, sediment and water plants was similar, supporting exchange of isolates among these niches (Fig. 4). Yet the water population was much richer in prevalence of the *STb* gene, and had no isolates with *hlyA*. Virulence gene distribution of snail populations was different, with more than half the isolates containing the *eaeA* gene. While all isolates from bovine feces belonged to phylogroup E, none contained any of the six virulence genes (Fig. 4). Few E isolates carried *Stx2*, and none tested positive for *Stx1* (Fig. S2). β-glucuronidase negative strains would not have formed green colonies on MLGA, and would have been excluded, so some phylogroup E strains containing virulence genes may have been excluded in our study.

**Fig. 4.**
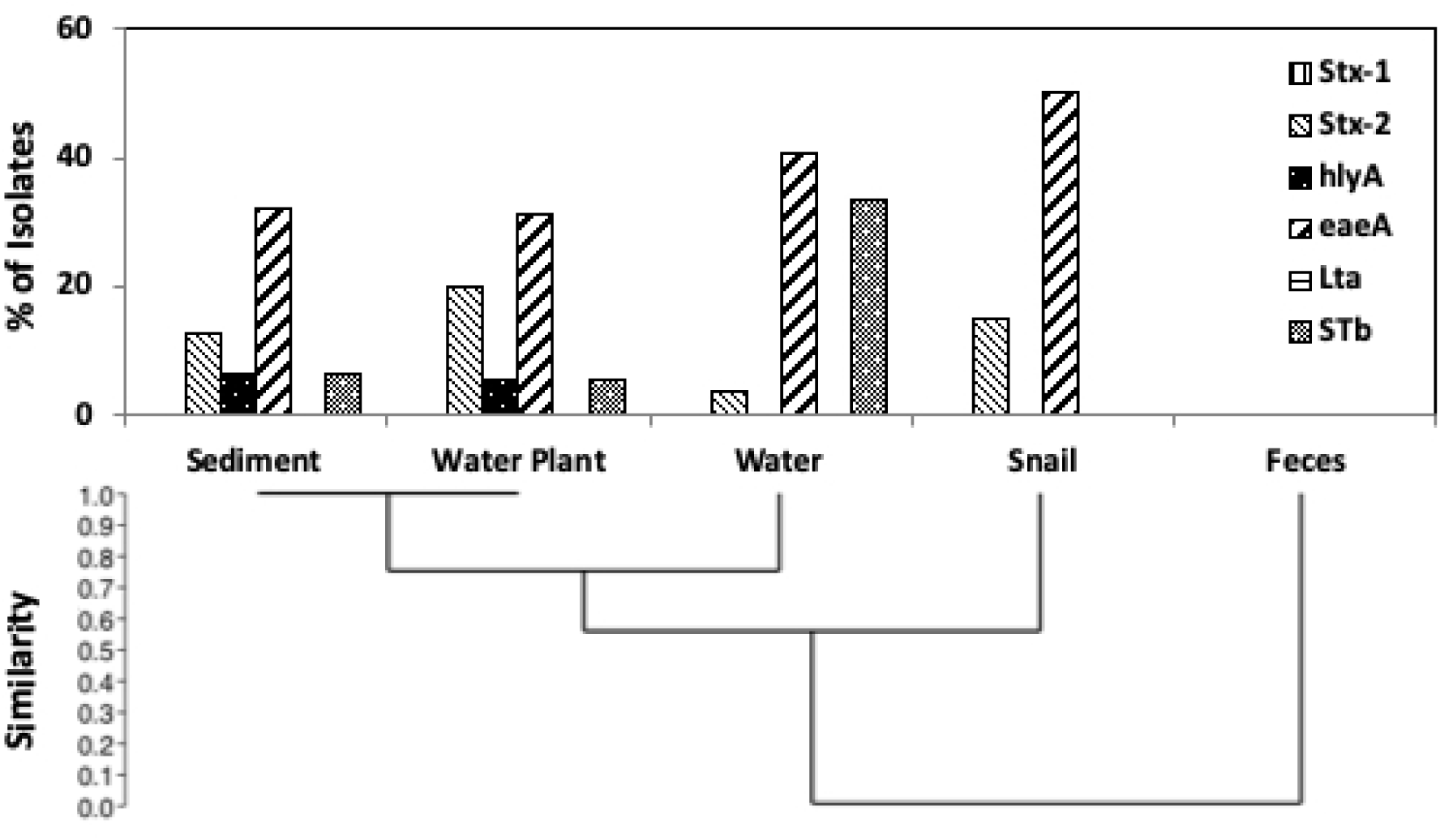
Virulence gene distribution across isolates from the five sample types. The relatedness between virulence gene distribution profiles was determined by UPGMA using the constrained Jaccard coefficient.

### Antibiotic Resistance profiling

One antibiotic from each of six target classes was chosen to evaluate the resistance of isolates: ceftriaxone (CRO, class cephalosporins), ciprofloxacin (CIP, class-fluoroquinolones), gentamicin (CN, class aminoglycosides), azithromycin (AZM, class-macrolides), meropenem (MEM, class carbapenems), and tetracycline (TE). Isolate collections from water and water plants showed a similar resistance distribution, with 60% of isolates resistant to gentamicin (Fig. 5). Sediment antibiotic resistance distribution was different from water and water plant populations. Water, water plant, and sediment samples contained isolates resistant to three antibiotics, many of which also contained the *eaeA* gene as well as either *STb* or *hlyA* (Fig. 6). The isolate collection from snails had a unique antibiotic resistance profile, with 80% sensitive to all antibiotics (Fig. 6), whereas only 20% of the isolates from water, sediment and water plant isolates were not resistant to any of the antibiotics. However, most of the snail isolates displayed intermediate resistance to three or four antibiotics (Fig. S3). Isolates from bovine feces also displayed a unique antibiotic resistance profile (Fig. 5), with 80% of isolates displaying intermediate resistance to either two or three antibiotics (Fig. S3).

**Fig. 5.**
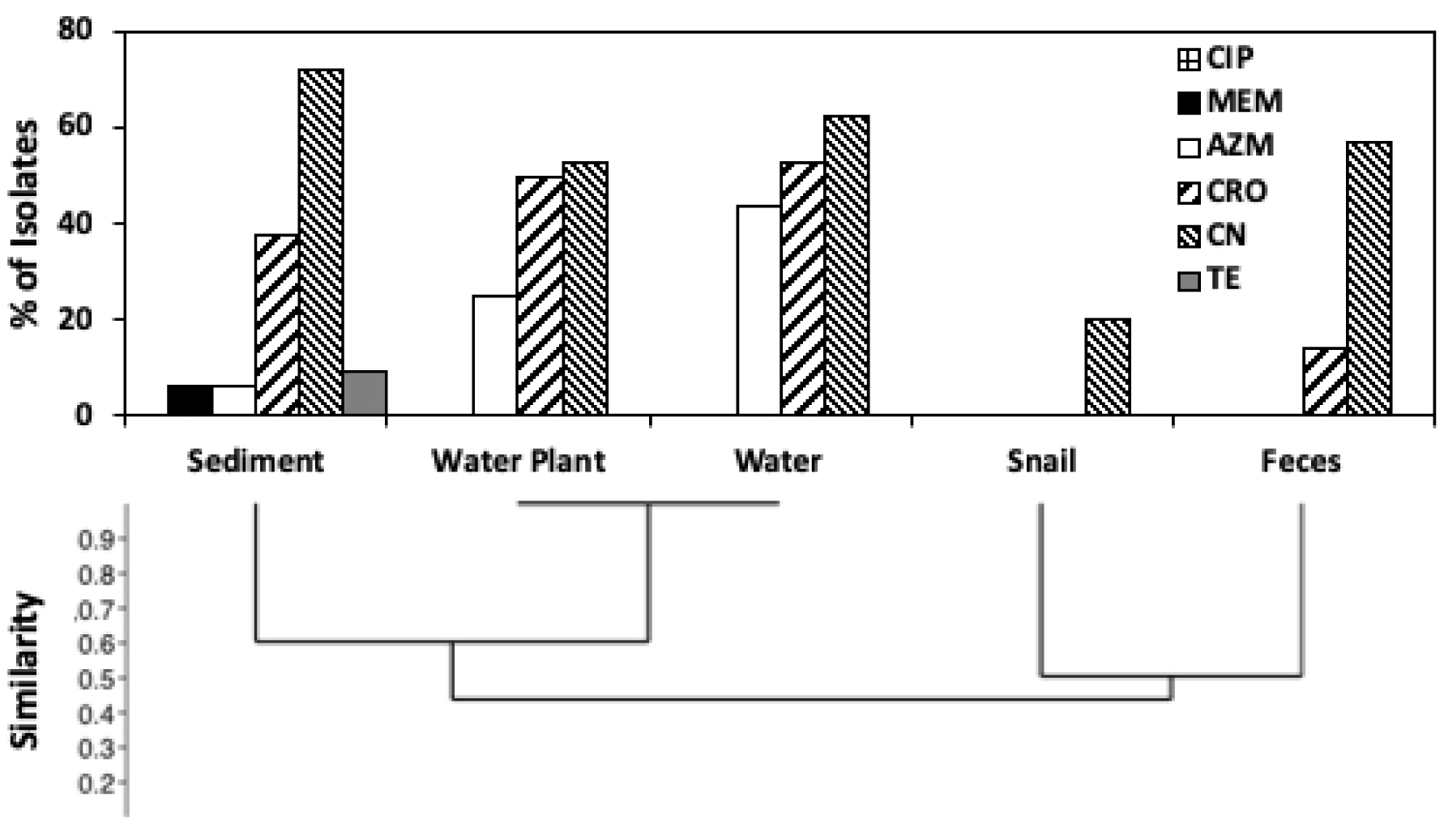
Antibiotic resistance across isolates from the five sample types. The relatedness between resistance profiles was determined by UPGMA using the constrained Jaccard coefficient.

**Fig. 6.**
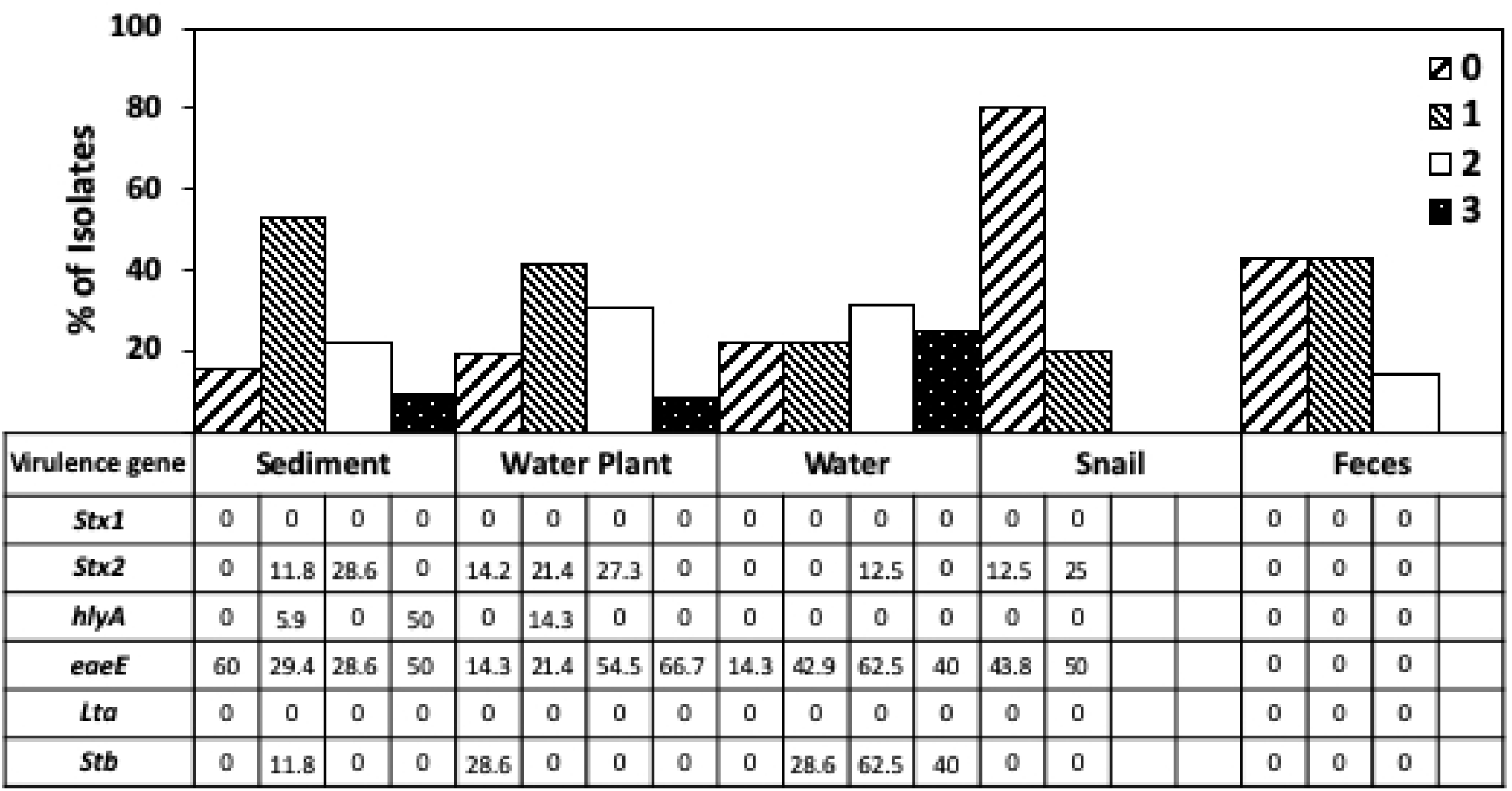
Sensitivity and multidrug resistance (resistance to 0, 1, 2 or 3 antibiotics) across sample types, compared to occurrence of virulence genes (percentage).

## Discussion

We sought to determine whether environmental *E. coli* display niche preference by associating with specific environments. We chose a secluded peri-urban pond adjacent to a cattle pasture, isolating *E. coli* from water, sediment, submerged water plants and water snails, as well as from freshly deposited bovine feces in the adjacent pasture. To obtain evidence of niche partitioning, isolates were characterized genotypically by phylogrouping, analysis of their *uidA* and *mutS* sequences, and virulence gene distribution, and phenotypically for antibiotic resistance.

Snail *E. coli* populations were predominated by phylogroup B2. Snail phylogroup distribution was different (p <0.001) to water, sediment and water plant polulations when using multinomial log-linear regression analysis. This indicated that strains display preference for either snail or aquatic niches but not both. Phylogroup distribution differed slightly between sediment and water and water plants but was not significant by multinomial log-linear regression analysis, indicating indiscrimate distribution of specific strains among these three niches. The prevalance of B1 and E, and some B2 in water, water plant and sediment was consistant with previous studies where B1 have been interpreted as generalists and harbor traits linked to plant association, whereas B2 strains are associated more with animals [53, 54]. Phylogroup distribution within the *E. coli* population in both water and superficial sediments showed spatial variation [31]. It has also been reported that phylogenetic groups are adaptable and genotypically influenced by changes in environmental conditions, however phylogroup B1 isolates seem to persist in water [8, 55]. Our data indicated that the B2 populations occurring in the pond persisted mostly in water snails. Likewise, phylogroup E strains predominated in bovine feces deposited nearby, and despite run-off from the pasture to the pond, did not thrive in the pond environment. The composition differences of phylogroups among populations in different environments may be caused by differences in adaptability and genome plasticity of *E. coli* strains [55]. Such variation in phylogroup distribution suggests that *E. coli* phylogroups are affected by niche specific selective pressures [54].

The phylogroup E strains isolated from the water, sediment and water plants formed several clusters within the *uidA mutS* phylogeny. Importantly, snail phylogroup E isolates clustered separately, as did bovine fecal isolates, indicating three separate groups of isolates and supporting niche preference among various phylogroup E strains. The *mutS* and *uidA* phylogeneny showed that some clusters were devoid of reference strains. None of our phylogroup E isolates clustered with any reference strains, suggesting these isolates are different to those typically associated with humans. In a recent study of cattle pasture we also found a higher percentage of phylogroup E in bovine fecal isolates compared to soil isolates, and none clustered with reference strains [9]. There appear to be diverse environmental β-Glucuronidase positive *E. coli* that are allocated to phylogroup E by the Clermont scheme [6], but that do not align with human isolates available in the databanks, warranting further investigation. The prevalence of *E. coli* in soils depends on specific conditions with phylogroup B1 and E associated with pasture lands while B2 and D phylogroups were associated with wooded areas [56]. Collectively, the phylogeny derived from *uidA* and *mutS* genes supported by phylogroup distribution analysis showed a preference of certain isolates with distinct backgrounds for specific niches.

Population genetic analysis of *mutS* and *uidA* supported the existence of three distinct genetic backgrounds within the collection of isolates analysed. The bovine fecal isolates had a homogenous background mostly lacking admixture. Some snail isolates shared this background, while others had their own backgound also lacking admixture. Isolates from water, water plants and sediment varied. Some had pure bovine background, others pure snail background, while the majority had an admixture of two or three backgrounds, bovine and / or snail plus a third, apparently aquatic one. This indicated directional gene flow from bovine fecal, and separately from snail-associated strains to aquatic strains. In contrast there was no or limited evidence for gene flow from aquatic to snail or cattle populations, indicating that none of these aquatic strains were able to persist in snail or bovine gastrointestinal environments. The neutral theory of molecular evolution makes a clear prediction on how the genetic drift in the absence of all other evolutionary forces shapes genetic diversity[57]. To study genomic evolution and consider a more complex explanation for the pattern of molecular variation, the neutral theory must be rejected as a null hypothesis [58]. Genetic variation in *E. coli* combines aspects of recombination, selection and population structure [59]. The gene flow model has some support in the literature. Retchless and Lawrence (60) proposed the fragment speciation model in which different segments of bacterial chromosomes become genetically isolated at different times. Sheppard, McCarthy (61) found evidence of increasing gene flow between previously distinct *Campylobacter* species. Luo, Walk (62) described the genomes of environmental isolates of *E.coli* and found little evidence of gene exchange between gut commensal *E. coli* due to possible ecological barriers, although they found transfer of core genes within the clades. Simlarly, Karberg, Olsen (63) found that recently acquired genes in *Salmonella* and *Escherichia* genomes have similar codon usage frequencies, while cores genes have noticeably diverged in codon usage. Therefores it seems that *Salmonella* and *Escherichia* strains acquire genes from common pangenomes shared among enterobacterial species.

The presence of virulence genes *eaeA, Stx2, hlyA*, and *STb* indicates potential pathogens, though it has been suggested that the occurrence of single or multiple virulence genes in *E. coli* does not confirm its pathogenicity, unless it has the appropriate combination of virulence genes to cause disease to the host. Enteric pathogens exposed to vegetables express similar genes to those required to colonize the host intestine, indicating that enteric bacteria may have the ability for colonization of vegetables by using similar mechanism required for animal cells [64]. High prevalence of the intimin-encoding gene *eaeA* was observed in all four pond niches, but not in feces, indicating presence of *eaeA* may play a role in aquatic fitness that is distinct from virulence. Byappanahalli, Nevers (65) detected a high level of *eaeA* in isolates from algae and to a lesser extent in those from water and sand samples from lake Michigan. *eaeA* is one of the most frequently detected *E. coli* pathogenicity genes in the environment [66-68]. It is not certain if these isolates with virulence genes are pathogenic and persist in the environment, or whether they acquire these genes from these environments.

The similarities in patterns of antibiotic resistance in aquatic populations suggested a common source of resistant strains, with preference for these niches. Snail populations were almost devoid of resistance to the wide aray on antibiotics evaluated, again supporting niche preference. *E. coli* isolated from various sampling sources showed variation in the antibiotic resistance patterns depending on the use of antibiotics and their exposure to environments [69-71]. This pond was not being used for any human or domestic animal activities and there was no direct input of wastewater. It is unclear whether isolates acquired antibiotic resistance through antibiotic exposure, or whether they maintain these genes in the absence of antibiotics [72].

In conclusion, sediment, water and water plant populations showed similarities in phylogroup distribution, occurrence of virulence genes and antibiotic resistance patterns in their populations, indicating that individual strains of this population can associate with any of these three niches. Snail-associated populations were different, and contained several apparently novel *E. coli* strains, primarily belonging to phylogroup B2. Bovine fecal populations from the adjoining pasture were different based on phenotype and genotype, and not similar to any aquatic isolates. The distinct distribution patterns of *E. coli* strains indicate niche preference, with specific aquatic strains not associating with snails or cattle.

## Acknowledgements

GN was supported by a fellowship from the South Dakota Agricultural Experiment Station.

